# Representation of auditory motion directions and sound source locations in the human planum temporale

**DOI:** 10.1101/302497

**Authors:** Ceren Battal, Mohamed Rezk, Stefania Mattioni, Jyothirmayi Vadlamudi, Olivier Collignon

**Affiliations:** Center of Mind/Brain Sciences, University of Trento, Italy; Institut de recherche en Psychologie (IPSY), Institute of Neuroscience (IoNS), Université catholique de Louvain (UcL)

**Keywords:** spatial hearing, planum temporale, auditory motion, auditory space, direction selectivity, fMRI, multivariate analyses

## Abstract

The ability to compute the location and direction of sounds is a crucial perceptual skill to efficiently interact with dynamic environments. How the human brain implements spatial hearing is however poorly understood. In our study, we used fMRI to characterize the brain activity of male and female humans listening to left, right, up and down moving as well as static sounds. Whole brain univariate results contrasting moving and static sounds varying in their location revealed a robust functional preference for auditory motion in bilateral human Planum Temporale (hPT). Using independently localized hPT, we show that this region contains information about auditory motion directions and, to a lesser extent, sound source locations. Moreover, hPT showed an axis of motion organization reminiscent of the functional organization of the middle-temporal cortex (hMT+/V5) for vision. Importantly, whereas motion direction and location rely on partially shared pattern geometries in hPT, as demonstrated by successful cross-condition decoding, the responses elicited by static and moving sounds were however significantly distinct. Altogether our results demonstrate that the hPT codes for auditory motion and location but that the underlying neural computation linked to motion processing is more reliable and partially distinct from the one supporting sound source location.

**SIGNIFICANCE STATEMENT:** In comparison to what we know about visual motion, little is known about how the brain implements spatial hearing. Our study reveals that motion directions and sound source locations can be reliably decoded in the human Planum Temporale (hPT) and that they rely on partially shared pattern geometries. Our study therefore sheds important new lights on how computing the location or direction of sounds are implemented in the human auditory cortex by showing that those two computations rely on partially shared neural codes. Furthermore, our results show that the neural representation of moving sounds in hPT follows a “preferred axis of motion” organization, reminiscent of the coding mechanisms typically observed in the occipital hMT+/V5 region for computing visual motion.

## INTRODUCTION

While the brain mechanisms underlying the processing of visual localization and visual motion have received considerable attention (Braddick et al., 2001; Movshon and Newsome, 1996; Newsome and Park, 1988), much less is known about how the brain implements spatial hearing. The representation of auditory space relies on the computations and comparison of intensity, temporal and spectral cues that arise at each ear (Blauert, 1982; Searle et al., 1976). In the auditory pathway, these cues are both processed and integrated in the brainstem, thalamus, and cortex in order to create an integrated neural representation of auditory space (Boudreau and Tsuchitani, 1968; Goldberg and Brown, 1969; Knudsen & Konishi 1978; Ingham et al., 2001; see Grothe et al., 2010 for review).

Similar to the dual-stream processing model in vision (Ungerleider and Mishkin, 1982; Goodale and Milner, 1992), partially distinct ventral “what” and dorsal “where” auditory processing streams have been proposed for auditory processing (Barrett and Hall, 2006; Lomber and Malhotra, 2008; Rauschecker and Tian, 2000; Recanzone, 2000; Romanski et al., 1999; Tian et al., 2001; Warren and Griffiths, 2003). In particular, the dorsal stream is thought to process sound source location and motion both in animals and humans (Arnott et al., 2004; Alain et al., 2001; Maeder et al., 2001; Rauschecker and Tian 2000; Tian et al., 2001). However, it remains poorly understood whether the human brain implements the processing of auditory motion and location using distinct, similar, or partially shared neural substrates (Poirier et al., 2017; Smith et al., 2010; Zatorre et al., 2002).

One candidate region that might integrate spatial cues to compute motion and location information is the planum temporale (hPT) (Barrett and Hall, 2006; Baumgart and Gaschler-Markefski, 1999; Warren et al., 2002). hPT is located in the superior temporal gyrus, posterior to Helsch’ gyrus, and is typically considered part of the dorsal auditory stream (Derey et al., 2016; Poirier et al., 2017; Rauschecker and Tian, 2000; Warren et al., 2002). Some authors have suggested that hPT engages in both the processing of moving sounds and the location of static sound-sources (Barrett and Hall, 2006; Derey et al., 2016; Krumbholz et al., 2005; Smith et al., 2004, 2007, 2010; Zatorre et al., 2002). This proposition is supported by early animal electrophysiological studies demonstrating the neurons in the auditory cortex that are selective to sound source location and motion directions (Altman, 1968, 1994; Benson et al., 1981; Doan et al., 1999; Imig et al., 1990; Middlebrooks and Pettigrew, 1981; Poirier et al., 1997; Rajan et al., 1990). In contrast, other studies in animals (Poirier et al., 2017) and humans (Baumgart and Gaschler-Markefski, 1999; Bremmer et al., 2001; Hall and Moore, 2003; Krumbholz et al., 2005; Lewis et al., 2000; Pavani et al., 2002; Poirier et al., 2005) pointed toward a more specific role of hPT for auditory motion processing.

The hPT, selective to process auditory motion/location in a dorsal “where” pathway is reminiscent of the dorsal region in the middle temporal cortex (hMT+/V5) in the visual system. hMT+/V5 is dedicated to process visual motion (Movshon and Newsome, 1996; Watson et al., 1993; Tootell et al., 1995) and displays a columnar organization in particular of a preferred axis of motion directions (Albright et al., 1984; Zimmermann et al., 2011). Whether hPT disclose similar characteristic tuning remains however unknown.

The main goals of the present study were threefold. First, using MVPA, we investigated whether information about auditory motion direction and sound-source location can be retrieved from the pattern of activity in hPT. Further, we asked whether the spatial distribution of the neural representation is in the format of “preferred axis of motion/location” as observed in the visual motion selective regions (Albright et al., 1984; Zimmermann et al., 2011). Finally, we aimed at characterizing whether the processing of motion direction (e.g. going to the left) and sound-source location (e.g. being in the left) rely on partially common neural representations in the hPT.

## MATERIALS AND METHODS

### Participants

Eighteen participants with no reported auditory problems were recruited for the study. Two participants were excluded due to poor spatial hearing performance in the task, as it was lower by more than 2.5 standard deviations than the average of the participants. The final sample included 16 right-handed participants (8 females, age range: 20 to 42, mean ± SD = 32 ± 5.7 years). Participants were blindfolded and instructed to keep their eyes closed throughout the experiments and practice runs. All the procedures were approved by the research ethics boards of the Centre for Mind/Brain Sciences (CIMeC) and University of Trento. Experiments were undertaken with the understanding and written consent of each participant.

### Auditory stimuli

Our limited knowledge of the auditory space processing in the human brain might be a consequence of the technical challenge of evoking vivid perceptual experience of auditory space while using neuroimaging tools such as fMRI, EEG, or MEG. In this experiment, to create an externalized ecological sensation of sound location and motion inside the MRI scanner, we relied on individual in-ear stereo recordings that were recorded in a semi-anechoic room and from 30 loudspeakers on horizontal and vertical planes, mounted on two semi-circular wooden structures with a radius of 1.1m (see Figure 1A). Participants were seated in the center of the apparatus with their head on a chin-rest, such that the speakers on the horizontal and vertical planes were equally distant from participants’ ears. Then, these recordings were re-played to the participants when they were inside the MRI scanner. By using such sound system with in-ear recordings, auditory stimuli automatically convolved with each individuals’ own pinna and head related transfer function to produce a salient auditory perception in external space.

**Figure 1.**
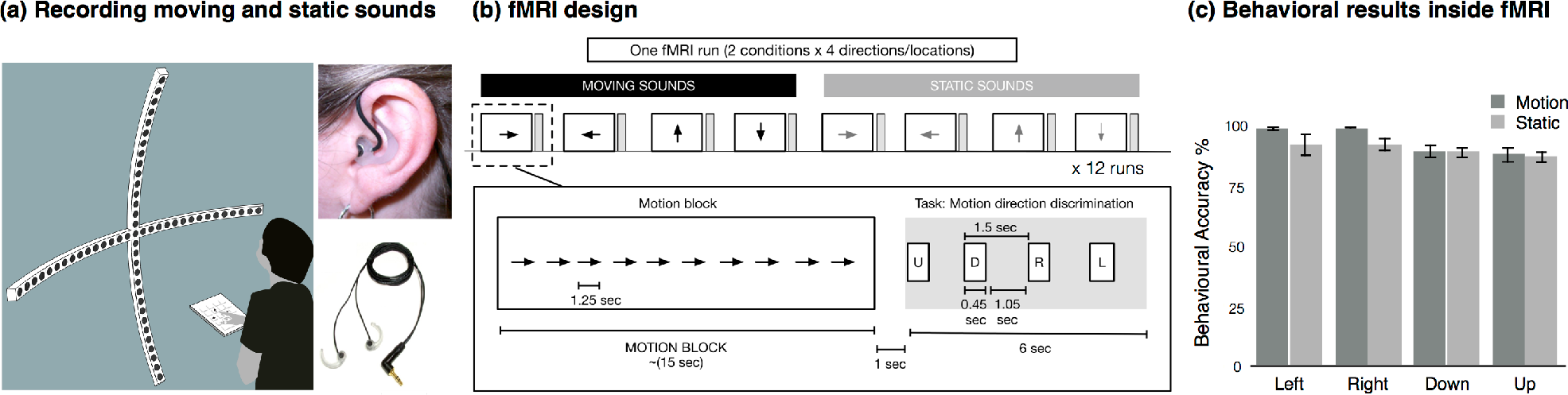
Stimuli and Experimental Design. **(A)**. The acoustic apparatus used to present auditory moving and static sounds while binaural recordings were carried out for each participant before the fMRI session. **(B)**. Auditory stimuli presented inside the MRI scanner consisted of 8 conditions: leftward, rightward, downward and upward moving stimuli and left, right, down and up static stimuli. Each condition was presented for 15 s (12 repetition of 1250 ms sound, no ISI) and followed by 7 s gap for indicating the corresponding direction/location in space and 8 s of silence (total inter-block interval was 15 s). Sound presentation and response button press were pseudo-randomized. Participants were asked to respond as accurately as possible during the gap period. **(C)**. The behavioural performance inside the scanner.

The auditory stimuli were prepared using custom MATLAB scripts (r2013b; Matworks). Auditory stimuli were recorded using binaural in-ear omni-directional microphones (Sound Professionals-TFB-2; ‘flat’ frequency range 20–20,000 Hz) connected to a portable Zoom H4n digital wave recorder (16-bit, stereo, 44.1 kHz sampling rate). Microphones were positioned at the opening of participant’s left and right auditory ear canals. While auditory stimuli were played, participants were listening without performing any task with their head fixed to the chin-rest in front of them. Binaural in-ear recordings allowed combining binaural properties such as interaural time and intensity differences, and participant specific monaural filtering cues to create reliable and ecological auditory space sensation (Pavani et al., 2002).

#### Stimuli recordings

Sound stimuli consisted of 1250 ms pink noise (50 ms rise/fall time). In the motion condition, the pink noise was presented moving in 4 directions: leftward, rightward, upward and downward. Moving stimuli covered 120° of space/visual field in horizontal and vertical axes. To create the perception of smooth motion, the 1250 ms of pink noise was fragmented into 15 equal length pieces with each 83.333 ms fragment being played every two speakers, and moved one speaker at a time, from outer left to outer right (rightward motion), or vice-versa for the leftward motion. For example, for the rightward sweep, sound was played through speakers located at −60° and −52° consecutively, followed by −44°, and so on. A similar design was used for the vertical axis. This resulted in participants perceiving moving sweeps covering an arc of 120° in 1250 ms (speed = 96°/s; 50 ms fade in/out) containing the same sounds for all four directions. The choice of the movement speed of the motion stimuli aimed to create listening experience relevant to everyday-life conditions. Moreover, at such velocity it has been demonstrated that human listeners are not able to make the differences between concatenated static stimuli from motion stimuli elicited by a single moving object (Poirier et al., 2017), supporting the participant’s report that our stimuli were perceived as smoothly moving (no perception of successive snapshots).

In the static condition, the same pink noise was presented separately at one of 4 locations: left, right, up, and down. Static sounds were presented at the second most outer speakers (−56° and +56° in the horizontal axis, and +56° and −56° in the vertical axis) in order to avoid possible reverberation difference at the outermost speakers. The static sounds were fixed at one location at a time instead of presented in multiple locations (Smith et al., 2008; 2010; Poirier et al., 2017; Krumbholz et al., 2005). This strategy was purposely adopted for two main reasons. First, randomly presented static sounds can evoke robust sensation of auditory apparent motion (Strybel & Neale 1994; Lakatos et al., 1997; see Carlile 2016 for review). Second, and crucially for the purpose of the present experiment, presenting static sounds located in a given position and moving sounds directed toward the same position allowed us to investigate whether moving and static sounds share a common representational format using cross-condition decoding (see below), which would have been impossible if the static sounds where randomly moving in space.

Before the recordings, the sound pressure levels (SPL) were measured from the participant’s head position and ensured that each speaker conveys 65dB-A SPL. All participants reported strong sensation of auditory motion and were able to detect locations with high accuracy (see Fig 1C). Throughout the experiment, participants were blindfolded. Stimuli recordings were conducted in a session that lasted approximately 10 minutes, requiring the participant to remain still during this period.

### Auditory experiment

Auditory stimuli were presented via MR-compatible closed-ear headphones (Serene Sound, Resonance Technology; 500-10KHz frequency response) that provided average ambient noise cancellation of about 30 dB-A. Sound amplitude was adjusted according to each participant’s comfort level. To familiarize the participants with the task, they completed a practice session outside of the scanner until they reached above 80% accuracy.

Each run consisted of the 8 auditory categories (4 motion and 4 static) randomly presented using a block-design. Each category of sound was presented for 15 s (12 repetition of 1250 ms sound, no ISI) and followed by 7 s gap for indicating the corresponding direction/location in space and 8 s of silence (total inter-block interval was 15 s). The ramp (50 ms fade in/out) applied at the beginning and at the end of each sound creates static bursts and minimized adaptation to the static sounds. During the response gap, participants heard a voice saying “left”, “right”, “up”, and “down” in pseudo-randomized order. Participants were asked to press a button with their right index finger when the auditory block’s direction or location was matching with the auditory cue (Figure 1B). The number of targets and the order (position 1-4) of the correct button press were balanced across category. This procedure was adopted to ensure that the participants gave their response using equal motor command for each category and to ensure the response is produced after the end of the stimulation period for each category. Participants were instructed to emphasize accuracy of response but not reaction times.

Each scan consisted of one block of each category, resulting in a total of 8 blocks per run, with each run lasting 4 m 10 s. Participants completed a total of 12 runs. The order of the blocks was randomized within each run, and across participants.

Based on pilot experiments, we decided to not rely on a sparse-sampling design as sometimes done in the auditory literature in order to present the sounds without the scanner background noise (Hall et al., 1999). These pilot experiments showed that the increase in the signal to noise ratio potentially provided by sparse sampling did not compensate for the loss in the number of volume acquisitions. Indeed, pilot recordings on participants not included in the current sample showed that, given a similar acquisition time between sparse-sampling designs (several options tested) and continuous acquisition, the activity maps elicited by our spatial sounds contained higher and more reliable beta values using continuous acquisition.

### fMRI data acquisition and analyses

#### Imaging parameters

Functional and structural data were acquired with a 4T Bruker MedSpec Biospin MR scanner, equipped with an 8-channel head coil. Functional images were acquired with T2*-weighted gradient echo-planar sequence. Acquisition parameters were: repetition time of 2500 ms, echo time of 26 ms, flip angle of 73°, a field of view of 192 mm, a matrix size of 64 × 64, and voxel size of 3 × 3 × 3 mm^3^. A total of 39 slices were acquired in ascending feet-to-head interleaved order with no gap. The three initial scans of each acquisition run were discarded to allow for steady-state magnetization. Before every two EPI runs, we performed an additional scan to measure the point-spread function (PSF) of the acquired sequence, including fat saturation, which served for distortion correction that is expected with high-field imaging (Zeng and Constable, 2002).

High-resolution anatomical scan was acquired for each participant using a T1-weighted 3D MP-RAGE sequence (176 sagittal slices, voxel size of 1×1×1 mm^3^; field of view 256 × 224 mm; repetition time = 2700 ms; TE = 4.18 ms; FA: 7°; inversion time: 1020 ms). Participants were blindfolded and instructed to lie still during acquisition and foam padding was used to minimize scanner noise and head movement.

#### Univariate fMRI analysis

##### Whole brain

Raw functional images were pre-processed and analysed with SPM8 (Welcome Trust Centre for Neuroimaging London, UK; http://www.fil.ion.ucl.ac.uk/spm/software/spm/) implemented in MATLAB R2014b (MathWorks). Before the statistical analysis, our preprocessing steps included slice time correction with reference to the middle temporal slice, realignment of functional time series, the coregistration of functional and anatomical data, spatial normalization to an echo planar imaging template conforming to the Montreal Neurological Institute space, and spatial smoothing (Gaussian kernel, 6 mm FWHM) were performed.

To obtain blood oxygen level-dependent (BOLD) activity related to auditory spatial processing, we computed single subject statistical comparisons with fixed-effect general linear model (GLM). In the GLM, we used eight regressors from each category (four motion direction, four sound source location). The canonical double-gamma hemodynamic response function implemented in SPM8 was convolved with a box-car function to model the above mentioned regressors. Motion parameters derived from realignment of the functional volumes (3 translational motion and 3 rotational motion parameters), button press, and the four auditory response cue events were modeled as regressors of no interest. During the model estimation, the data were high-pass filtered with cut-off 128s to remove the slow drifts/ low-frequency fluctuations from the time series. To account for serial correlation due to noise in fMRI signal, autoregressive (AR (1)) was used.

In order to obtain activity related to auditory processing in the whole brain, the contrasts tested the main effect of each category ([Left Motion], [Right Motion], [Up Motion], [Down Motion], [Left Static], [Right Static], [Up Static], [Down Static]). To find brain regions responding preferentially to the moving and static sounds, we combined all motion conditions [Motion] and all static categories [Static]. The contrasts tested the main effect of each condition ([Motion], [Static]), and comparison between the conditions ([Motion > Static], and [Static > Motion]). These linear contrasts generated statistical parametric maps (SPM[T]) which were further spatially smoothed (Gaussian kernel 8 mm FWHM) and entered in a second-level analysis, corresponding to a random effects model, accounting for inter-subject variance. One-sample t-tests were run to characterize the main effect of each condition ([Motion], [Static]), and the main effect of motion processing ([Motion > Static]) and static location processing ([Static > Motion]). Statistical inferences were performed at a threshold of *p*<0.05 corrected for multiple comparisons (Family-Wise Error corrected; FWE) either over the entire brain volume or after correction for multiple comparisons over small spherical volumes (12 mm radius) located in regions of interest. Significant clusters were anatomically labeled using the xjView Matlab toolbox (http://www.alivelearn.net/xjview) or structural neuroanatomy information provided in the Anatomy Toolbox (Eickhoff et al., 2007).

#### Region of interest analysis

##### ROI Definition

Due to the hypothesis-driven nature of our study we defined hPT as an *a priori* region of interest for statistical comparisons and in order to define the volume in which we performed multivariate pattern classification analyses.

To avoid any form of double dipping that may arise when defining the ROI based on our own data, we decided to independently define hPT, using a meta-analysis method of quantitative association test, implemented via the online tool Neurosynth (Yarkoni et al., 2011) using the term query “Planum Temporale”. Rather than showing which regions are disproportionately reported by studies where a certain term is dominant (uniformity test; P (activation | term)), this method identifies regions whose report in a neuroimaging study is diagnostic of a certain term being dominant in the study (association test; P (term | activation)). As such, the definition of this ROI was based on a set of 85 neuroimaging studies at the moment of the query (September 2017). This method provides an independent method to obtain masks for further region-of-interest analysis. The peak coordinate from the meta-analysis map used to create a 6 mm spheres (117 voxels) around the peak z-values of hPT (peak MNI coordinates [-56 −28 8] and [60 −28 8]; lhPT and rhPT hereafter, respectively).

Additionally, hPT regions of interest were individually defined using anatomical parcellation with FreeSurfer (http://surfer.nmr.mgh.harvard.edu). The individual anatomical scan was used to perform cortical anatomical segmentation according to the Destrieux (Destrieux et al., 2010) atlas. We selected planum temporale label defined as [G_temp_sup-Plan_tempo, 36] bilaterally. We equated the size of ROI across participants and across hemispheres to 110 voxels (each voxel being 3mm isotropic). For anatomically defined hPT ROIs, all further analyses were carried out in subject space for enhanced anatomical precision and to avoid spatial normalization across participants. We replicated our pattern of results in anatomically defined parcels (left and right hPT) obtained from the single subject brain segmentation (for further analysis see Battal et al., 2018).

#### ROI Analyses

##### Univariate

The beta parameter estimates of the 4 motion directions and 4 sound source locations were extracted from lhPT and rhPT regions (Fig 2C). In order to investigate the presence of motion directions/sound source locations selectivity and condition effect in hPT regions, we performed a 2 Conditions (hereafter refers to the stimuli type: motion, static) × 4 Orientations (hereafter refers to either direction or location of the stimuli: left, right, down, and up) repeated measures ANOVA in each hemisphere separately on these beta parameter estimates. Statistical results were then corrected for multiple comparisons (number of ROIs x number of tests) using the false discovery rate (FDR) method (Benjamini and Yekutieli, 2001). A Greenhouse–Geisser correction was applied to the degrees of freedom and significance levels whenever an assumption of sphericity was violated.

#### ROI - Multivariate pattern analyses

##### Within Condition Classification

Four-class and binary classification analyses were conducted within the hPT region in order to investigate the presence of auditory motion direction and sound source location information in this area. To perform multi-voxel pattern classification, we used a univariate one-way ANOVA to select a subset of voxels (n=110) that are showing the most significant signal variation between the categories of stimuli (in our study, between orientations). This feature-selection not only ensures a similar number of voxels within a given region across participants (dimensionality reduction), but, more importantly, identifies and selects voxels that are carrying the most relevant information across categories of stimuli (Cox and Savoy 2003; De Martino et al., 2008), therefore minimizing the chance to include in our analyses voxels carrying noises unrelated to our categories of stimuli.

Multivariate pattern analyses (MVPA) were performed in the lhPT and rhPT. Preprocessing steps were identical to the steps performed for univariate analyses, except for functional time series that were smoothed with a Gaussian kernel of 2 mm (FWHM). MVPA was performed using CoSMoMVPA (http://www.cosmomvpa.org/; (Oosterhof et al., 2016), implemented in MATLAB. Classification analyses were performed using support vector machine (SVM) classifiers as implemented in LIBSVM (http://www.csie.ntu.edu.tw/~cjlin/libsvm; Chang and Lin, 2011). A general linear model was implemented in SPM8, where each block was defined as a regressor of interest. A beta map was calculated for each block separately. Two multi-class and six binary linear SVM classifiers with a linear kernel with a fixed regularization parameter of C = 1 were trained and tested for each participant separately. The two multi-class classifiers were trained and tested to discriminate between the response patterns of the 4 auditory motion directions and locations, respectively. Four binary classifiers were used to discriminate brain activity patterns for motion and location within axes (left vs. right motion, left vs. right static, up vs. down motion, up vs. down static, hereafter within axis classification). We used 8 additional classifiers to discriminate across axes (Left vs. Up, Left vs. Down, Right vs. Up, and Right vs. Down motion directions, Left vs. Up, Left vs. Down, Right vs. Up, and Right vs. Down sound source locations, hereafter across axes classification).

For each participant, the classifier was trained using a cross-validation leave-one-out procedure where training was performed with n-1 runs and testing was then applied to the remaining one run. In each cross-validation fold, the beta maps in the training set were normalized (z-scored) across conditions, and the estimated parameters were applied to the test set. To evaluate the performance of the classifier and its generalization across all the data, the previous step was repeated 12 times where in each fold a different run was used as the testing data and the classifier was trained on the other 11 runs. For each region per participant, a single classification accuracy was obtained by averaging the accuracies of all cross-validation folds.

#### Cross-condition classification

To test whether motion directions and sound source locations share a similar neural representation in hPT region, we performed cross-condition classification. We carried out the same steps as for the within-condition classification as described above but trained the classifier on sound source locations and tested on motion directions, and vice versa. The accuracies from the two cross-condition classification analyses were averaged. For interpretability reasons, cross-condition classification was only interpreted on the stimuli categories that the classifiers discriminated reliably (above chance level) for both motion and static conditions (e.g. if discrimination of left vs. right was not successful in one condition, either static or motion, then the left vs. right cross-condition classification analysis was not carried out).

#### Within-orientation classification

To foreshadow our results, cross-condition classification analyses (see previous section) showed that motion directions and sound source locations share, at least partially, a similar neural representation in hPT region. To further investigate the similarities/differences between the neural patterns evoked by motion directions and sound source locations in the hPT, we performed 4 binary classifications in which the classifiers were trained and tested on the same orientation pairs: leftward motion vs. left static, rightward motion vs. right static, upward motion vs. up static, and downward motion vs. down static. If the same orientation (leftward and left location) across conditions (motion and static) generates similar patterns of activity in hPT region, the classifier would not be able to differentiate leftward motion direction from left sound location. However, significant within-orientation classification would indicate that the evoked patterns within hPT contain differential information for motion direction and sound source location in the same orientation (e.g. left).

The mean of the four binary classifications was computed to produce one accuracy score per ROI. Prior to performing the within-orientation and cross-condition MVPA, each individual pattern was normalised separately across voxels so that any cross or within-orientation classification could not be due to global univariate activation differences across the conditions.

#### Statistical analysis: MVPA

Statistical significance in the multivariate classification analyses was assessed using non-parametric tests permuting condition labels and bootstrapping (Stelzer et al., 2013). Each permutation step included shuffling of the condition labels and re-running the classification 100 times on the single-subject level. Next, we applied bootstrapping procedure in order to obtain a group-level null distribution that is representative of whole group. From each individual’s null distribution one value was randomly chosen and averaged across all the participants. This step was repeated 100,000 times resulting in a group level null distribution of 100,000 values. The classification accuracies across participants was considered as significant if p<0.05 after corrections for multiple comparisons (number of ROIs x number of tests) using the FDR method (Benjamini and Yekutieli, 2001).

Similar approach was adopted to assess statistical difference between the classification accuracies of two auditory categories (e.g. four motion direction vs. four sound source location, left motion vs. left static, left motion vs. up motion, etc.). We performed additional permutation tests (100,000 iterations) by building a null distribution for t-stats after randomly shuffling the classification accuracy values across two auditory categories and re-calculating the two-tail t-test between the classification accuracies of these two categories. All p-values were corrected for multiple comparisons (number of ROIs x number of tests) using the FDR method (Benjamini and Yekutieli, 2001).

#### Representation Similarity analysis

##### Neural Dissimilarity matrices

To further explore the differences in the representational format between sound source locations and motion directions in hPT region, we relied on representation similarity analysis (RSA; Kriegeskorte et al., 2008). More specifically, we tested the correlation between the representational dissimilarity matrix (RDM) of right and left hPT in each participant with different computational models that included condition-invariant models assuming orientation invariance across conditions (motion, static), condition-distinct models assuming that sound source location and motion direction sounds elicit highly dissimilar activity patterns, and a series of intermediate graded models between them. The RSA was performed using CosmoMVPA toolbox (Oosterhof et al., 2016) implemented in MATLAB. To perform this analysis, we first extracted in each participant the activity patterns associated with each condition (Edelman et al., 1998; Haxby et al., 2001). Then, we averaged individual subject statistical maps (i.e. activity patterns) in order to have a mean pattern of activity for each auditory category across runs. Finally, we used Pearson’s linear correlation as the similarity measure to compare each possible pair of the activity patterns evoked by the four different motion directions and four different sound source locations. This resulted in an 8 × 8 correlation matrix for each participant that was then converted into a representational dissimilarity matrix (RDMs) by computing 1 – correlation. Each element of the RDM contains the dissimilarity index between the patterns of activity generated by two categories, in other words the RDM represents how different is the neural representation of each category from the neural representations of all the other categories in the selected ROI. The 16 neural RDMs (1 per participant) for each of the 2 ROIs were used as neural input for RSA.

#### Computational models

To investigate shared representations between auditory motion directions and sound source locations, we created multiple computational models ranging from a fully condition-distinct model to a fully condition-invariant model with intermediate gradients in between (Zabicki et al., 2016) (Figure 4C).

#### Condition-Distinct model

The condition-distinct models assume that dissimilarities between motion and static condition is 1 (i.e. highly dissimilar), meaning that neural responses/patterns generated by motion and static conditions are totally unrelated. For instance, there would be no similarity between any motion directions with any sound source location. The dissimilarity values in the diagonal were set to 0, simply reflecting that neural responses for the same direction/location are identical to themselves.

#### Condition-Invariant model

The condition-invariant models assume a fully shared representation for specific/corresponding static and motion conditions. For example, the models consider the neural representation for the left sound source location and the left motion direction highly similar. All within-condition (left, right, up and down) comparisons are set to 0 (i.e. highly similar) regardless of their auditory condition. The dissimilarities between different directions/locations are set to 1 meaning that each within condition sound (motion or static) is different from all the other within conditions.

#### Intermediate models

To detect the degree of similarity/shared representation between motion direction and sound source location patterns, we additionally tested 2 classes of 5 different intermediate models. The two classes were used to deepen the understanding of characteristic tuning of hPT for separate direction/location or axis of motion/location. The two model classes represent 2 different possibilities. The first scenario was labeled as Within-Axis Distinct, and these models assume that each of the 4 directions/locations (i.e. left, right, up, down) would generate a distinctive neural representation different from all of the other within-condition sounds (e.g. the patterns of activity produced by the left category are highly different from the patterns produced by right, up and down categories) (see Figure 4C, upper panel). To foreshadow our results, we observed preference for axis of motion in MVP-classification, therefore we created another class of models to further investigate neural representations of within-axis and across-axes of auditory motion/space. The second scenario was labeled with Within-Axis Combined, and these models assume that opposite direction/locations within the same axis would generate similar patterns of activity (e.g. the pattern of activity of horizontal (left and right) categories are different from the patterns of activity of vertical categories (up and down) (see Figure 4C, lower panel).

In all intermediate models, the values corresponding to the dissimilarity between same auditory spaces (e.g. left motion and left location) were gradually modified from 0.9 (motion and static conditions are mostly distinct) to 0.1 (motion and static conditions mostly similar). These models were labeled IM9, 7, 5, 3, and 1 respectively.

In all condition-distinct and intermediate models, the dissimilarity of within-condition sounds was fixed to 0.5 and dissimilarity of within-orientation sounds was fixed to 1. Across all models, the diagonal values were set to 0.

#### Performing RSA

We computed Pearson’s correlation to compare neural RDMs and computational model RDMs. The resulting correlation captures which computational model better explains the neural dissimilarity patterns between motion direction and sound source location conditions. To compare computational models, we performed Mann-Whitney-Wilcoxon rank-sum test between for every pair of models. All p-values were then corrected for multiple comparisons using the FDR method (Benjamini and Yekutieli, 2001). To test differences between two classes of models (Within-Axis Combined vs. Within-Axis Distinct), within each class, the correlation coefficient values were averaged across hemispheres and across models. Next, we performed permutation tests (100,000 iterations) by building a null distribution for differences between classes of models after randomly shuffling the correlation coefficient values across two classes, and re-calculating the subtraction between the correlation coefficients of Within-Axis Combined and Within-Axis Distinct classes.

To visualize the distance between the patterns of the motion directions and sound source locations, we used multi-dimensional scaling (MDS) to project the high-dimensional RDM space onto 2 dimensions with the neural RDMs that were obtained from both lhPT and rhPT.

Additionally, the single-subject 8 × 8 correlation matrices were used to calculate the reliability of the data considering the signal-to-noise ratio of the data (Kriegeskorte et al., 2007). For each participant and each ROI, the RDM was correlated with the averaged RDM of the rest of the group. The correlation values were then averaged across participants. This provided the maximum correlation that can be expected from the data.

## RESULTS

### Behavioral results

During the experiment, we collected target direction/location discrimination responses (see Figure 1C). The overall accuracy scores were entered into 2 × 4 (Condition, Orientation) repeated measures ANOVA. No main effect of Condition (F_1,15_ = 2.22; p = 0.16) was observed, indicating that the overall accuracy while detecting direction of motion or sound source location did not differ. There was a significant main effect of orientation (F_1.6,23.7_ = 11.688; p = 0.001), caused by greater accuracy in the horizontal orientations (left and right) as compared to the vertical orientations (up and down). Post-hoc two-tailed t-tests (Bonferroni corrected) revealed that accuracies did not reveal significant difference within horizontal orientations (left vs right; t_15_ = −0.15, p=1), and within vertical orientations (up vs down; t_15_ = 0.89, p = 1). However, left orientation accuracy was greater as compared to down (t_15_ = 3.613, p = 0.005), and up (t_15_ = 4.51, p < 0.001) orientations and right orientation accuracy was greater as compared to the down (t_15_ = 3.76, p = 0.003) and up (t_15_ = 4.66, p < 0.001) orientation accuracies. No interaction between Condition × Orientation was observed, pointing out that differences between orientations in terms of performance expresses both for static and motion. A Greenhouse–-Geisser correction was applied to the degrees of freedom and significance levels whenever an assumption of sphericity was violated.

### fMRI results – whole-brain univariate analyses

To identify brain regions that are preferentially recruited for auditory motion processing, we performed a univariate RFX-GLM contrast [Motion > Static] (Figure 2B). Consistent with previous studies (Dormal et al., 2016; Getzmann and Lewald, 2012; Pavani et al., 2002; Poirier et al., 2005; Warren et al., 2002), whole-brain univariate analysis revealed activation in the superior temporal gyri, bilateral hPT, precentral gyri, and anterior portion of middle temporal gyrus in both hemispheres (Figure 2B, Table 1). The most robust activation (resisting whole brain FWE correction, p<0.05) was observed in the bilateral hPT (peak MNI coordinates [−46 −32 10] and [60 −32 12]). We also observed significant activation in occipito-temporal regions (in the vicinity of anterior hMT+/V5) as suggested by previous studies (Dormal et al., 2016; Poirier et al., 2005; Warren et al., 2002).

**Table 1.** Results of the univariate analyses for the main effect of auditory motion processing [motion > static], and auditory localization processing [static > motion]. Coordinates reported in this table are significant (p < 0.05 FWE) after correction over small spherical volumes (SVC, 12 mm radius) of interest (#) or over the whole brain (*). Coordinates used for correction over small spherical volumes are as follows (x, y, z, in MNI space): left middle temporal gyrus (hMT+/V5) [-42 −64 4] (Dormal et al., 2016), right middle temporal gyrus (hMT +/V5) [42 − 60 4] (Dormal et al., 2016), right superior frontal sulcus [32 0 48] (Collignon et al., 2011), right middle occipital gyrus [48 −76 6] (Collignon et al., 2011). K represents the number of voxels when displayed at p(unc) < 0.001. L: left, R: right, G: gyrus, S: sulcus.

| Area | k | x<br>(mm) | y<br>(mm) | z<br>(mm) | Z | p |
| --- | --- | --- | --- | --- | --- | --- |
| *MOTION > STATIC* |  |  |  |  |  |  |
| L planum temporale | 10506 | -46 | -32 | 10 | 6.63 | 0.000* |
| L Middle Temporal G |  | -56 | -38 | 14 | 6.10 | 0.000* |
| L Precentral G |  | -46 | -4 | 52 | 5.25 | 0.004* |
| L Putamen |  | -22 | 2 | 2 | 4.98 | 0.011* |
| L Middle Temporal G | 43 | -50 | -52 | 8 | 3.79 | 0.01# |
| R Superior Temporal G | 7074 | 66 | -36 | 12 | 6.44 | 0.000* |
| R Superior Temporal G |  | 62 | -2 | -4 | 5.73 | 0.000* |
| R Superior Temporal G |  | 52 | -14 | 0 | 5.56 | 0.001* |
| R Precentral G |  | 50 | 2 | 50 | 4.70 | 0.032* |
| R Superior Frontal S | 159 | 46 | 0 | 50 | 4.40 | 0.001# |
| R Middle Temporal G | 136 | 42 | -60 | 6 | 4.31 | 0.001# |
| R Middle Occipital G | 24 | 44 | -62 | 6 | 3.97 | 0.006# |

### fMRI results – ROI univariate analyses

Beta parameter estimates were extracted from the pre-defined ROIs (see methods) for the four motion directions and four sound source locations from the auditory experiment (Figure 2C). We investigated the condition effect and the presence of direction/location selectivity in lhPT and rhPT regions separately by performing 2 × 4 (Conditions, Orientations) repeated measures of ANOVA with beta parameter estimates. In lhPT, main effect of Conditions was significant (F_1,15_ = 37.28, p < 0.001), indicating that auditory motion evoked higher response compared to static sounds. There was no significant main effect of Orientations (F_1.5,22.5_ = 0.77, p = 0.44), and no interaction (F_3,45_ = 2.21, p = 0.11). Similarly, in rhPT, only main effect of Conditions was significant (F_1,15_ = 37.02, p < 0.001). No main effect of Orientation (F_1.5,23.2_ = 1.43, p = 0.26) or interaction (F_3,45_ = 1.74, p = 0.19) was observed. Overall, brain activity in the hPT as measured with beta parameter estimate extracted from univariate analysis did not provide evidence of motion direction or sound source location selectivity.

**Figure 2.**
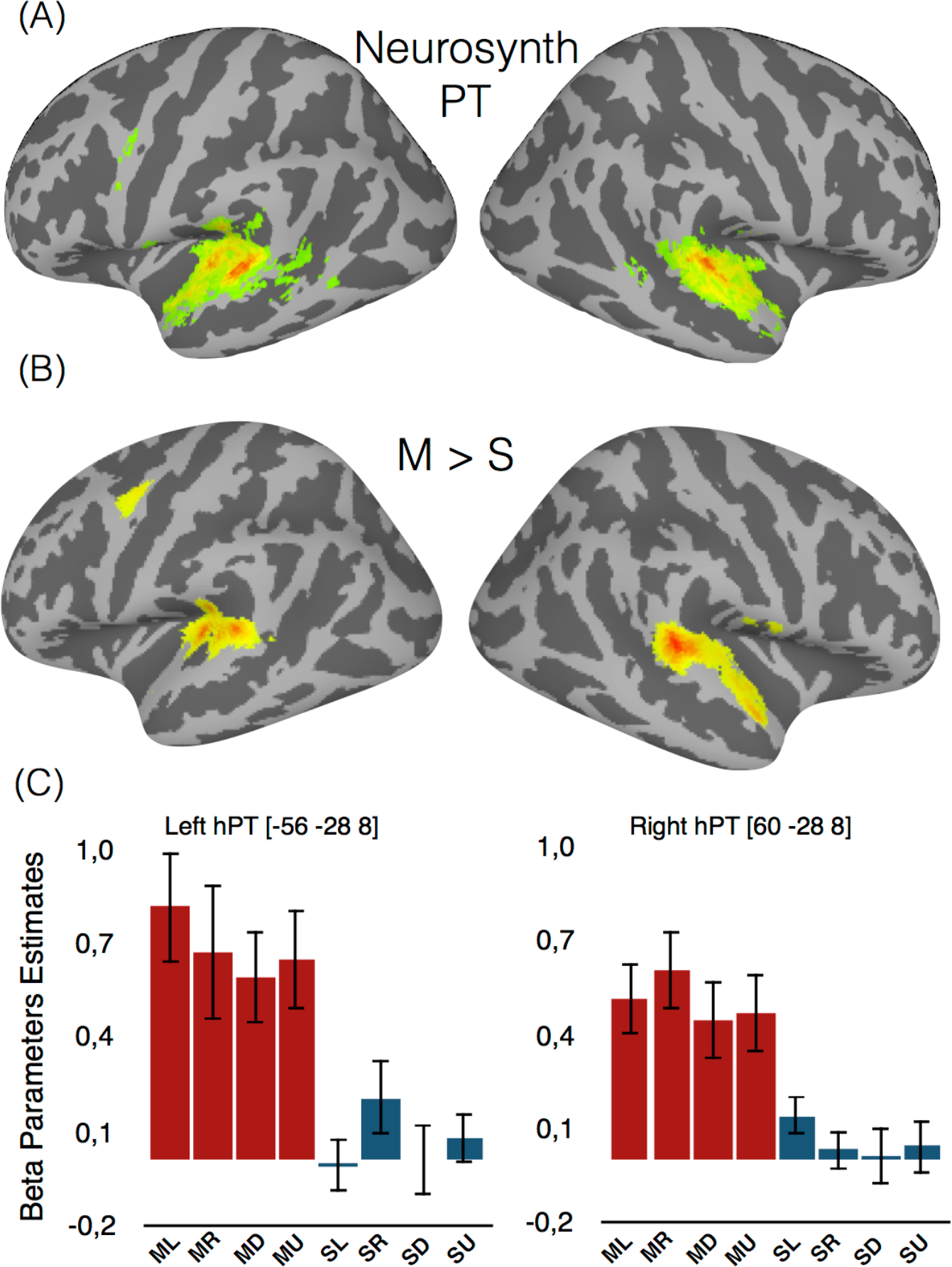
Univariate whole brain results. **(A)**. Association test map was obtained from the online tool Neurosynth using the term “Planum Temporale” (FDR corrected p <05). The black spheres are illustration of drawn mask (radius = 6mm, 117 voxels) around the peak coordinate from Neurosynth (search term “planum temporale”, meta-analysis of 85 studies). **(B)**. Auditory motion processing [motion > static] thresholded at p <0.05 FWE-corrected. **(C)**. Mean activity estimates (arbitrary units ± SEM) associated with the perception of auditory motion direction (red) and sound-source location (blue). ML: motion left, MR: motion right, MD: motion down, MU: motion up, SL: static left, SR: static right, SD: static down, and SU: static up.

### fMRI results – ROI multivariate pattern analyses

To further investigate the presence of information about auditory motion direction and sound source location in hPT, we ran multi-class and binary multivariate pattern classification. Figure 3A-C shows the mean classification accuracy across categories in each ROI.

### MVPA – Within Condition

Multi-class across four conditions classification accuracy in the hPT was significantly above chance (chance level = 25%) in both hemispheres for motion direction (lhPT: mean ± SD = 38.4 ±7, p < 0.001; rhPT: mean ± SD = 37.1 ± 6.5, p<0.001), and sound source location (lhPT: mean ± SD = 32.4 ± 6.7, p < 0.001; rhPT: mean ± SD = 31.2 ± 7.5, p < 0.001) (Figure 3A). In addition, we assessed the differences between classification accuracies for motion and static stimuli by using permutation tests in lhPT (p = 0.024) and rhPT (p = 0.024), indicating greater accuracies for classifying motion direction than sound source location across all regions.

### MVPA – Binary Within-Axis

Binary horizontal (left vs. right) within-axis classification showed significant results in both lhPT and rhPT for static sounds (lhPT: mean ± SD = 58.6 ± 14.5, p<0.001; rhPT: mean ± SD = 56.5 ± 11.9, p = 0.008) (Figure 3C), while motion classification was significant only in the rhPT (mean ± SD = 55.5 ± 13.9, p = 0.018) (Figure 3B). Moreover, binary vertical (up vs down) within-axis classification was significant only in the lhPT for both motion (mean ± SD = 55.7 ± 7.1, p = 0.01), and static (mean ± SD = 54.9 ± 11.9, p = 0.03) conditions (Figure 3B-C).

### MVPA – Binary Across-Axis

We used 8 additional binary classifiers to discriminate across axes moving and static sounds. Binary across-axes (Left vs. Up, Left vs. Down, Right vs. Up, and Right vs. Down) classifications were significantly above chance level in the bilateral hPT for motion directions (Left vs. Up lhPT: mean ± SD = 65.8 ±14.8, p<0.001; rhPT: mean ± SD = 64.8 ± 9.4, p<0.001; Left vs. Down lhPT: mean ± SD = 74 ±15.9, p<0.001; rhPT: mean ± SD = 66.9 ± 10.4, p<0.001; Right vs. Up lhPT: mean ± SD = 72.4 ±15.8, p<0.001; rhPT: mean ± SD = 71.4 ± 13.3, p<0.001; Right vs. Down lhPT: mean ± SD = 73.4 ±11.8, p<0.001; rhPT: mean ± SD = 68.2 ± 15.9, p<0.001) (Figure 3B). However, sound source locations across-axes classifications were significant only in certain conditions (Left vs. Up rhPT: mean ± SD = 56 ± 12.7, p=0.018; Left vs. Down lhPT: mean ± SD = 55.2 ±11.2, p=0.024; rhPT: mean ± SD = 57.3 ± 17.2, p=0.003; Right vs. Down lhPT: mean ± SD = 57.6 ±8.4, p=0.005) (Figure 3C).

### MVPA – “Axis of Motion” Preference

In order to test whether neural patterns within hPT contain information about opposite directions/locations within an axis, we performed four binary within-axis classifications. Similar multivariate analyses were performed to investigate the presence of information about across-axes directions/locations. The classification accuracies were plotted in Figure 3B-C.

In motion direction classifications, to assess the statistical difference between classification accuracies of across axes (left vs. up, left vs. down, and right vs. up, right vs. down) and within axes (left vs. right, and up vs. down) directions, we performed pairwise permutation tests and FDR-corrected for multiple comparisons. Across-axes classification accuracies in lhPT ([left vs. up] vs. [left vs. right]: p=0.006, [left vs. down] vs. [left vs. right]: p<0.001, [right vs. down] vs. [left vs. right]: p<0.001, [right vs. up] vs. [left vs. right]: p=0.001), and rhPT ([left vs. up] vs. [left vs. right]: p=0.029, [left vs. down] vs. [left vs. right]: p=0.014, [right vs. down] vs. [left vs. right]: p=0.02, [right vs. up] vs. [left vs. right]: p=0.003) were significantly higher compared to the horizontal within-axis classification accuracies. Similarly, across-axes classification accuracies were significantly higher when compared with vertical within-axis classification accuracies in lhPT ([up vs. down] vs. [left vs. up], p=0.02; [up vs. down] vs. [left vs. down], p=0.001; [up vs. down] vs. [right vs. up], p=0.001; [up vs. down] vs. [right vs. down], p <0.001) and rhPT ([up vs. down] vs. [left vs. up], p= 0.001; [up vs. down] vs. [left vs. down], p=0.001; [up vs. down] vs. [right vs. up], p= 0.001; [up vs. down] vs. [right vs. down], p=0.002). No significant difference was observed between the within-axis classifications in lhPT ([left vs. right] vs. [up vs. down], p=0.24) and rhPT ([left vs. right] vs. [up vs. down], p=0.31). Similarly, no significance difference was observed among the across-axes classification accuracies in the bilateral hPT.

In static sound location classifications, no significant difference was observed between across-axes and within-axes classification accuracies, indicating that the classifiers did not perform better when discriminating sound source locations across axes compared to the opposite locations.

**Figure 3.**
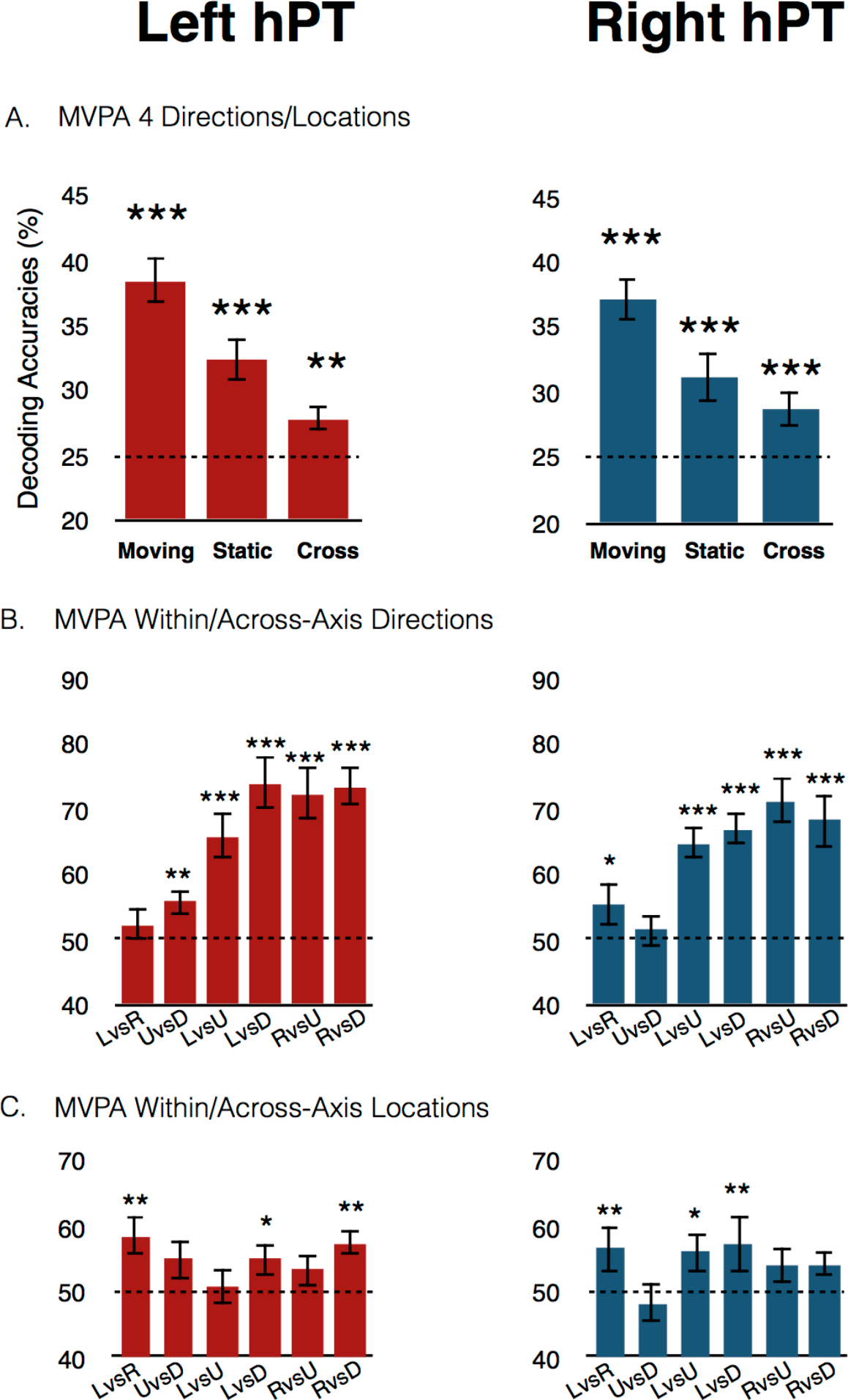
Within and cross-classification results. **(A)**. Classification results for the 4 conditions in functionally defined hPT region. Within-condition and cross-condition classification results are shown in the same bar plots. Moving: four motion direction; Static: four sound source location; and Cross: cross-condition classification accuracies. **(B)**. Classification results of within (left vs. right, up vs. down) and across axes (left vs. up, left vs. down, right vs. up, right vs. down) motion directions. **(C)**. Classification results of within (left vs. right, up vs. down) and across axes (left vs. up, left vs. down, right vs. up, right vs. down) sound source locations. LvsR: Left vs. Right, UvsD: Up vs. Down, LvsU: Left vs. Up, LvsD: Left vs. Down, RvsU: Right vs. Up, RvsD: Right vs. Down classifications. FDR corrected p-values: (*) p<0.05, (**) p<0.01, (***) p<0.001 testing differences against chance level (dotted lines; see methods).

One may wonder whether the higher classification accuracy for across compared to within axes relates to the perceptual differences (e.g. difficulty in localizing) in discriminating sounds within the horizontal and vertical axes. Indeed, because we opted for an ecological design reflecting daily-life listening condition, we observed, as expected, that discriminating vertical directions was more difficult than discriminating horizontal ones (Middlebrooks and Green, 1991). In order to address this issue, we replicated our MVP-classification analyses after omitting the four participants showing the lowest performance in discriminating the vertical motion direction, leading to comparable performance (at the group level) within and across axes. We replicated our pattern of results by showing preserved higher classification accuracies across-axes than within-axis. Moreover, while accuracy differences between across- and within-axes classification was only observed in the motion condition, behavioral differences were observed in both static and motion conditions. To assess whether the higher across axes classification accuracies are due to difference in difficulties between horizontal and vertical sounds, we performed correlation analysis. For each participant, the behavioral performance difference between horizontal (left and right) and vertical (up and down) conditions was calculated. The difference between the classification accuracies within axes (left vs. right and up vs. down) and across axes (left vs. up, left vs. down, right vs. up, and right vs. down) was calculated. Spearman correlation between the (**Δ**) behavioral performance and (**Δ**) classification accuracies was not significant in the bilateral hPT (lhPT *r* = 0.18, p = 0.49; rhPT *r* = 0.4, p = 0.12). An absence of correlation suggests that there is no relation between the difference in classification accuracies and the perceptual difference. These results strengthen the notion that “axis of motion” organization observed in PT does not simply stem from behavioral performance differences.

### MVPA – Cross-condition

To investigate if motion direction and sound source locations rely on shared representation in hPT, we trained the classifier to distinguish neural patterns from the motion directions (e.g. going to the left) and then tested on the patterns elicited by static conditions (e.g. being in the left), and vice versa.

Cross-condition classification revealed significant results across 4 direction/location (lhPT: mean ± SD = 27.8 ± 5.3, p = 0.008, rhPT: mean ± SD = 28.7 ± 3.8, p<0.001) (Figure 3A). Within-axis categories did not reveal any significant cross-condition classification. These results suggest that a partial overlap between the neural patterns of moving and static stimuli in the hPT.

### MVPA – Within-orientation

Cross-condition classification results indicated a shared representation between motion directions and sound source locations. Previous studies argued that successful cross-condition classification reflects an abstract representation of stimuli conditions (Fairhall and Caramazza, 2013; Higgins et al., 2017; Hong et al., 2012). To test this hypothesis, patterns of the same orientation of motion and static conditions (e.g. leftward motion and left location) were involved in within-orientation MVPA. The rational was that if the hPT region carries fully abstract representation of space, within-orientation classification would give results in favor of the null hypothesis (no differences within the same orientation). In the within-orientation classification analysis, accuracies from the four within-orientation classification analyses were averaged and survived FDR corrections in bilateral hPT (lhPT: mean ± SD = 65.6 ± 5, p < 0.001, rhPT: mean ± SD = 61.9 ± 5.6, p<0.001), indicating that the neural patterns of motion direction can be reliably differentiated from sound-source location within hPT (Figure 4A).

**Figure 4.**
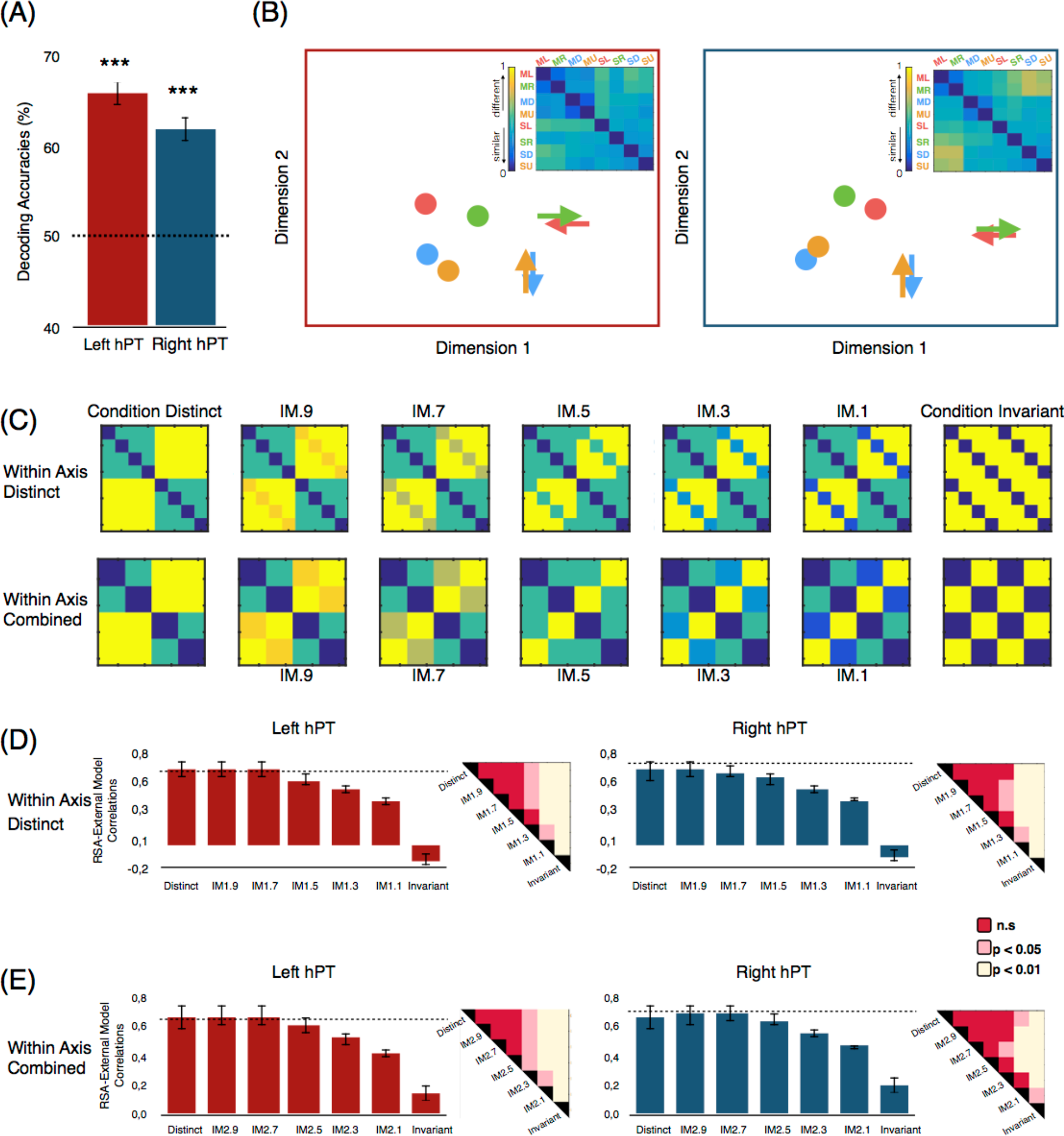
Pattern dissimilarity between motion directions and sound source locations. **(A)**. Across-condition classification results of across 4 conditions are represented in each ROI (lhPT and rhPT). 4 binary classifications [leftward motion vs. left location], [rightward motion vs. right location], [upward motion vs. up location], and [downward motion vs. down location] were computed and averaged to produce one accuracy score per ROI. FDR corrected p-values: (***) p<0.001. Dotted lines represent chance level. **(B)**. The embedded top panel shows neural RDMs extracted from left and right hPT, and multi-dimensional scaling (MDS) plot visualizes the similarities of the neural pattern elicited by 4 motion directions (arrows) and 4 sound source locations (dots). Color codes for arrow/dots are as follows: green indicates left direction/location, red indicates right direction/location, orange indicates up direction/location, and blue indicates down direction/location. ML: motion left, MR: motion right, MD: motion down, MU: motion up, SL: static left, SR: static right, SD: static down, and SU: static up. **(C-E)**. The results of representational similarity analysis (RSA) in hPT are represented. **(C)**. RDMs of the computational models that assume different similarities of the neural pattern based on auditory motion and static conditions. **(D-E)**. RSA results for every model and each ROI. For each ROI, the dotted lines represent the reliability of the data considering the signal-to-noise ratio (see Materials and Methods), which provides an estimate of the highest correlation we can expect in a given ROI when correlating computational models and neural RDMs. Error bars indicate SEM. IM1: Intermediate models with within-axis conditions distinct, IM2: Intermediate model with within-axis conditions combined. Each right up corner of the bar plots show visualization of significant differences for each class of models and hemispheres separately (Mann-Whitney-Wilcoxon rank-sum test, FDR corrected).

### RSA

#### Multi-dimensional Scaling

Visualization of the representational distance between the neural patterns evoked by motion directions and sound source locations further supported that within-axis directions show similar geometry compared to the across-axes directions, therefore, it is more difficult to differentiate the neural patterns of opposite directions in MVP-classification. MDS also showed that in both lhPT and rhPT, motion directions and sound source locations are separated into 2 clusters (Figure 4B).

### RSA with external models

The correlation between model predictions and neural RDMs for the lhPT and rhPT is shown in Figure 4D. The cross-condition classification results indicated a shared representation within the neural patterns of hPT for motion and static sounds. We examined the correspondence between the response pattern dissimilarities elicited by our stimuli with 14 different model RDMs that included a fully condition distinct, fully condition-invariant models, and intermediate models with different degrees of shared representation.

First class of computational RDMs were modeled with the assumption that the neural patterns of within-axis sounds are fully distinct. The analysis revealed a negative correlation with the fully condition-invariant model in the bilateral hPT (lhPT: mean r ± SD = −0.12 ± 0.18, rhPT: mean r ± SD = −0.01 ± 0.2) that increased gradually as the models progressed in the condition-distinct direction. The model that best fit the data was the M9 model in the bilateral hPT (lhPT: mean r ± SD = 0.66± 0.3, rhPT: mean r ± SD = 0,65 ± 0.3). A similar trend was observed for the second class of models that have the assumption of within-axis sounds evoke similar neural patterns. Condition-invariant model provided the least explanation of the data (lhPT: mean r ± SD = 0.14 ± 0.25, rhPT: mean r ± SD = 0.2 ± 0.29), and correlations gradually increased as the models progressed in the condition-distinct direction. The winner models in this class were the models M9 in lhPT and M7 in the rhPT (lhPT: mean r ± SD = 0.67 ± 0.22, rhPT: mean r ± SD = 0.69 ± 0.15).

In addition, we assessed differences between correlation coefficients for computational models using Mann-Whitney-Wilcoxon rank-sum test for each class of models and hemispheres separately (Figure 4D-E). In Within-Axis Distinct class in lhPT, pairwise comparisons of correlation coefficients did not show significant difference for [Condition Distinct vs. IM.9; p = 0.8], [Condition Distinct vs. IM7; p = 0.6; rhPT], [Condition Distinct vs. IM.5; p = 0.09], however as the models progressed in further condition invariant direction, the difference between correlation coefficients for models reached significance ([Condition Distinct vs. IM.3], p = 0.012; [Condition Distinct vs. IM.1], p = 0.007; [Condition Distinct vs. Condition Invariant], p <0.001), indicating Condition Distinct model provided stronger correlation compared to the models representing conditions similarly. In rhPT, the rank-sum tests between each pairs revealed no significant difference for [Condition Distinct vs. IM.9; p = 0.9], [Condition Distinct vs. IM7; p = 0.7], [Condition Distinct vs. IM.5; p = 0.3], and also [Condition Distinct vs. IM.3; p = 0.06], however as the models progressed in further condition invariant direction, the difference between correlation coefficients for models reached significance ([Condition Distinct vs. IM.1], p =0.006; [Condition Distinct vs. Condition Invariant], p <0.001).

The two classes of models were used to deepen our understanding of characteristic tuning of hPT for separate direction/location or axis of motion/location. To reveal the differences between Within-Axis Distinct and Within-Axis Combine classes of models, we relied on 2-sided signed-rank test. Within-Axis Combined models explained our stimuli space better than Within-Axis Distinct models supporting similar pattern representation within planes (p = 0.0023).

## DISCUSSION

In line with previous studies, our univariate results demonstrate higher activity for moving than for static sounds in the superior temporal gyri, bilateral hPT, precentral gyri, and anterior portion of middle temporal gyrus in both hemispheres (Baumgart and Gaschler-Markefski, 1999; Krumbholz et al., 2005; Pavani et al., 2002; Poirier et al., 2005; Warren et al., 2002). The most robust cluster of activity was observed in the bilateral hPT (Figure 2B, Table 1). Moreover, activity estimates extracted from independently defined hPT also revealed higher activity for moving relative to static sounds in this region. Both whole-brain and ROI analyses therefore clearly indicated a functional preference (expressed here as higher activity level estimates) for motion processing over sound-source location in bilateral hPT regions (Figure 2).

Does hPT contain information about specific motion directions and sound source locations? At the univariate level, our four (left, right, up and down) motion directions and sound source locations did not evoke differential univariate activity in hPT region (see Figure 2C). We then carried out multivariate pattern classification and observed that bilateral hPT contains reliable distributed information about the four auditory motion directions (Figure 3). Our results are therefore similar to the observations made with fMRI in the human visual motion area hMT+/V5 showing reliable direction-selective information despite comparable voxel-wise univariate activity levels across directions (Kamitani and Tong, 2006; but see Zimmermann et al., 2011). Within-axis MVP-classification results revealed that even if both horizontal (left versus right), and vertical (up versus down) motion directions can be classified in the hPT region (see Figure 3C-D), across axes (e.g. left versus up) direction classifications revealed much higher decoding accuracies compared to within-axis classifications. Such enhanced classification accuracy across axes versus within axis is reminiscent of observations made in MT+/V5 where a large-scale axis of motion selective organization was observed in non-human primates (Albright et al., 1984), and in human area MT+/V5 (Zimmermann et al., 2011). Further examination with RSA provided additional evidence that within-axis combined models (aggregating the opposite directions/location) explain better the neural representational space of hPT by showing higher correlations values compared to within-axis distinct models (see Figure 4D-E). Our findings suggest that the topographic organization principle of hMT+/V5 and hPT shows similarities in representing motion directions. Animal studies have shown that hMT+/V5 contains motion direction selective columns with specific directions organized side by side with their respective opposing motion direction counterparts (Albright et al., 1984; Born and Bradley, 2005; Diogo et al., 2003; Geesaman et al., 1997), an organization also probed using fMRI in humans (Zimmermann et al., 2011; but see below for alternative accounts). The observed difference for within versus between axis direction decoding may potentially stem from such underlying cortical columnar organization (Bartels et al., 2008; Haynes and Rees, 2006; Kamitani and Tong, 2005). Alternatively, it has been suggested that classifying orientation preference reflects much larger scale (e.g. retinotopy) rather than columnar organization (Op de Beeck, 2010; Freeman et al., 2011, 2013). Interestingly, high-field fMRI studies showed that the fMRI signal carries information related to both large- and fine-scale (columnar level) biases (Gardumi et al., 2016; Pratte et al., 2016; Sengupta et al., 2017). The present study sheds important new lights on the coding mechanism of motion direction within the hPT and demonstrates that fMRI signal in the hPT contains direction specific information and point toward an “axis of motion” organization. This result highlights intriguing similarities between the visual and auditory systems, arguing for common neural coding strategies of motion directions across the senses. However, further studies are needed to test the similarities between the coding mechanisms implemented in visual and auditory motion selective regions, and more particularly, what drives the directional information captured with fMRI in the auditory cortex.

Supporting univariate motion selectivity results in bilateral hPT, MVPA revealed higher classification accuracies for moving than for static sounds (Figure 3A-B). However, despite minimal univariate activity elicited by sound-source location in hPT, and the absence of reliable univariate differences in the activity elicited by each position (see Figure 2C), within-axis MVP-classification results showed that beside the vertical axis (up versus down), sound source location information can be reliably decoded bilaterally in hPT (Figure 3C). Our results are in line with previous studies showing that posterior regions in auditory cortex exhibit location sensitivity both in animals (Recanzone, 2000; Stecker et al., 2005; Tian et al., 2001) and humans (Ahveninen et al., 2006, 2013; Brunetti et al., 2005; Deouell et al., 2007; Derey et al., 2016; Krumbholz et al., 2005; Warren and Griffiths, 2003; Zatorre et al., 2002; Alain et al., 2001). In contrast to what was observed for motion direction however, within-axis classifications did not reveal lower decoding accuracies when compared to across-axis classifications in hPT for static sounds. This indicates that auditory sound source locations might not follow similar topographic representations than the one observed for motion directions.

To which extend the neural representation of motion directions and sound source locations overlaps has long been debated (Grantham, 1986; Kaas et al., 1999; Poirier et al., 2017; Romanski et al., 2000; Smith et al., 2004, 2007; Zatorre and Belin, 2001). A number of neurophysiological studies have reported direction-selective neurons along the auditory pathway (Ahissar et al. 1992; Spitzer and Semple 1993; Doan et al. 1999; Stumpf et al. 1992; Toronchuk et al. 1992; McAlpine et al. 2000). However, whether these neurons exhibit selectivity to motion direction specifically, or also to spatial location remains debated (Poirier et al. 1997; McAlpine et al. 2000; Ingham et al. 2001; Oliver 2003). In our study, we revisit this question by relying on cross-condition classification that revealed that auditory motion (e.g. going to the left) and sound-source location (being on the left) share partial neural representations in hPT (Figure 3A). This observation suggests that there is a shared level of computation between sounds located on a given position and sounds moving towards this position. Low-level features of these two types of auditory stimuli vary in many ways and produce large difference at the univariate level in the cortex (see Figure 2B). However, perceiving, for instance, a sound going toward the left side or located on the left side evoke a sensation of location/direction in the external space that is common across conditions. Our significant cross-condition classification may therefore relate to the evoked sensation/perception of an object being/going to a common external spatial location. Electrophysiological studies in animals demonstrated that motion-selective neurons in the auditory cortex displayed similar response profile to sounds located or moving toward the same position in external space, suggesting that the processing of sound-source locations may contribute to the perception of moving sounds (Ahissar et al., 1992; Doan et al., 1999; Poirier et al., 1997). Results from human psychophysiological and auditory evoked potential studies also strengthen the notion that sound source location contributes to motion perception (Getzmann and Lewald, 2011; Strybel and Neale, 1994). Our cross-condition MVPA results therefore extend the notion that motion directions and sound source locations might have common features that are partially shared for encoding spatial sounds.

Significant cross-condition classification based on multivariate fMRI signal has typically been considered as a demonstration that the region implements a partly common and abstracted representation of the tested conditions (Fairhall and Caramazza, 2013; Higgins et al., 2017; Hong et al., 2012). For instance, a recent study elegantly demonstrated that the human auditory cortex at least partly integrates interaural time and level differences (ITD and ILD) into a higher-order representation of auditory space based on significant cross-cue classification (training on ITD and classifying ILD, and reversely) (Higgins et al., 2017). In the present study, we argue that even if cross-condition MVP-classification indeed demonstrate the presence of partially shared information across sound source location and direction; successful cross-MVPA results should however not be taken as evidence that the region implements a purely abstract representation of auditory space. Our successful across-condition classification (see Figure 4A) demonstrated that, even though there are shared representations for moving and static sounds within hPT, classifiers are able to easily distinguish motion directions from sound source locations (e.g. leftward versus left location). RSA analyses further supported the idea that moving and static sounds elicit distinct patterns in hPT (see Figure 4B-D). Altogether, our results suggest that hPT contains both motion direction and sound-source location information but that the neural patterns related to these two conditions are only partially overlapping. Our observation of significant cross-condition classification based on highly distinct pattern of activity between static and moving sounds may support the notion that even if location information could serve as a substrate for movement detection, motion encoding does not solely rely on location information (Ducommun et al., 2002; Getzmann, 2011; Poirier et al., 2017).

## Acknowledgements

We would also like to express our gratitude to Giorgia Bertonati, Marco Barilari, Stefania Benetti, Valeria Occelli, Stephanie Cattoir who helped with the data acquisition; Jorge Jovicich for helping setting-up the fMRI acquisition parameters and Pietro Chiesa for continuing support with the auditory hardware. The project was funded by the ERC starting grant MADVIS (Project: 337573) awarded to OC. MR and JV are PhD fellows and OC is research associate at the Fond National de la Recherche Scientifique de Belgique (FRS-FNRS).

